# Divergence in photoperiod responses of a classical biological control agent *Neogalerucella calmariensis* (Coleoptera: Chrysomelidae) across a climatic and latitudinal gradient

**DOI:** 10.1101/2020.04.13.039974

**Authors:** Tyson Wepprich, Fritzi S. Grevstad

## Abstract

A key knowledge gap in classical biological control is to what extent insect agents evolve to novel environments. The introduction of biological control agents to new photoperiod regimes and climates may disrupt the coordination of diapause timing that evolved to the growing season length in the native range. We tested whether populations of *Neogalerucella calmariensis* (L.) have evolved in response to the potential mismatch of their diapause timing since their intentional introduction to the United States from Germany in the 1990s. Populations collected from 39.4° to 48.8° latitude in the western USA were reared in growth chambers to isolate the effects of photoperiod on diapause induction and development time. For all populations, shorter day lengths increased the proportion of beetles that entered diapause instead of reproducing. The critical photoperiods, or the day length at which half of a population diapauses, differed significantly among the sampled populations, generally decreasing at lower latitudes. The latitudinal trend reflects changes in growing season length, which determines the number of generations possible, and in local day lengths at the time beetles are sensitive to this cue. Development times were similar across populations, with one exception, and did not vary with photoperiod. These results show that there was sufficient genetic variation from the two German source populations to evolve different photoperiod responses across a range of environmental conditions. This study adds to the examples of rapid evolution of seasonal adaptations in introduced insects.

## Introduction

A key concern in classical biological control is whether insect agents evolve to novel environmental conditions and how this should inform introduction strategies in the new range (McEvoy et al. 2012, Roderick et al. 2012). Evolution may be expected in traits regulating insect lifecycles as local adaptations to the growing season length and environmental cues in the native range are disrupted (Saikkonen et al. 2012, Grevstad and Coop 2015). Multivoltine insects, which could potentially fit more than one generation in a year, have evolved genetic-based responses to environmental cues that determine when populations switch developmental pathways from reproduction to diapause in advance of winter (Danilevskii 1961, Tauber et al. 1986, Denlinger et al. 2017). Day length, which varies seasonally and across latitudes, is the principal cue that insects in temperate zones use, in conjunction with other cues from maternal effects, weather, or host plants, to initiate diapause development (Tauber et al. 1986, Bradshaw and Holzapfel 2007). The response typically takes the form of a switch from reproductive to diapause states when a sensitive life stage is exposed to a day length that is shorter than a threshold value. This is defined as the critical day length or critical photoperiod, used interchangeably in the literature to describe an individual’s threshold or the median threshold in a population.

The ubiquity of latitudinal clines in the critical photoperiod that induces diapause in native and long-established insects (reviewed by Masaki 1961, Danilevskii 1965, Tauber et al. 1986) suggests that similar selective pressures will drive geographic divergence in these seasonal adaptations in introduced insects. For two main reasons, the critical photoperiod is expected to evolve to be longer at higher latitudes: (1) summer day lengths used as cues are longer at higher latitudes and (2) shorter growing seasons at higher latitudes mean that insects need to induce diapause earlier in the summer when days are longer to prepare for an earlier winter onset. While day length for a given date is a strict function of latitude, growing season length is more locally variable depending on its climate.

In this study, we experimentally compared the diapause response and development rates in seven populations of *Neogalerucella calmariensis* (L.) (formerly *Galerucella calmariensis* L.), a leaf beetle introduced across North America as a biological control agent for the wetland weed purple loosestrife (*Lythrum salicaria* L.). Our study is preceded by a phenology model applied to this insect (Grevstad and Coop 2015) that demonstrates how introductions to new photoperiod and climate regimes may disrupt the coordination of diapause timing that evolved to the growing season length and photoperiod regime at the population’s origin. Newly introduced insects can experience mismatches between the number of generations *attempted* based on photoperiod-induced diapause and the *potential* number of generations that could be possible with a well-adapted photoperiodic response (Bean et al. 2007, Grevstad and Coop 2015). Attempting reproduction too late in the year could lead to high mortality as life stages unable to diapause confront winter or senescing host plants (Van Dyck et al. 2015, Kerr et al. 2020). Entering diapause too early in the growing season can lead to greater overwintering mortality because of the added duration of diapause over the summer (Bean et al. 2007). Other adaptations might include faster development to fit full generations within the available growing season or accelerated development in response to shortened day lengths toward the end of the season (McEvoy et al. 2012, Lindestad et al. 2019). The mismatches between season length and thermal time requirements to complete the final generation set up selective pressure for the photoperiod response, development time, and voltinism to adapt to novel climates and day lengths at different latitudes.

Biological control introductions draw from limited source populations in the agent’s native range so that prerelease host range tests accurately assess the risk of nontarget attacks (Hinz et al. 2014). Constrained genetic variation may limit the success of agents released across a wide range of environmental conditions (Roderick et al. 2012, Wright and Bennett 2018). While often assumed that biological control agents undergo a period of adaptation following introduction, documented examples are limited to a few cases (Wright and Bennett 2018). They include changes in springtime phenology in the ragwort flea beetle *Longitarsus jacobaeae* (Szucs et al. 2012), divergence in developmental rates between alpine and valley populations of the cinnabar moth *Tyria jacobaeae* (McEvoy et al. 2012), and a shortening of the critical photoperiod that cues diapause in the tamarisk leaf beetle *Diorhabda carinulata* (Bean et al. 2012). In the case of the *D. carinulata*, the evolution in photoperiod response allowed populations to increase voltinism and extend geographic range boundaries southward into regions where the original photoperiod response initially prevented establishment (Bean et al. 2007, 2012).

For *N. calmariensis*, we predicted that photoperiod responses had evolved to correspond with season length and day length cues across a latitudinal gradient, based on the potential for mismatched diapause timing modeled in Grevstad and Coop (2015). In this study, we reared beetles in controlled environmental chambers set with different artificial day lengths to test for genetic differences in photoperiod thresholds for diapause induction and photoperiod-dependent development rates. We show that, within 27 years post-introduction, these two traits have diverged across populations in a manner consistent with local adaptation.

## Materials and Methods

### Study system

*Neogalerucella calmariensis* was first introduced in 1992 across the United States and Canada as a classical biological control agent for purple loosestrife (*L. salicaria*), a noxious invasive weed common in wetlands (Hight et al. 1995). It established populations at sites with different climate regimes and photoperiod exposure (36-53°N) compared to its two source populations in Germany (50 and 54°N) (Hight et al. 1995, Corrigan et al. 2013). *Neogalerucella calmariensis* was introduced alongside its congener *Neogalerucella pusilla* (Duftschmidt), which coexists in the same habitat, or even on the same plant, without apparent differences in life history or ecological niche (Blossey 1995). We focus on *N. calmariensis* in this study, although *N. pusilla* was also present at some of the sites. We identified *N. calmariensis* in the field and later in the lab by their larger size, dark stripes on their elytra, and orange-brown coloration (Manguin et al. 1993).

*Neogalerucella calmariensis* has a facultative multivoltine lifecycle in North America. Adults overwinter and emerge as reproductive regardless of photoperiod exposure in spring (Blossey 1995, Bartelt et al. 2008). Based on degree-day (base 10°C) requirements, oviposition from overwintered adults occurs around 100 accumulated degree-days and subsequent development from egg to adult to oviposition requires around 523 degree-days (McAvoy and Kok 2004, Grevstad and Coop 2015). Newly eclosed (teneral) adults from subsequent generations are photoperiod sensitive and will enter diapause in response to short days (Velarde et al. 2002) and non-reproductive adults will start oviposition at a delay if exposed to longer days (Grevstad 1999). The baseline critical photoperiod (=critical day length) in beetles introduced from Germany was not measured experimentally. However, soon after release, it was estimated as <15.2 day hours by Bartelt et al. (2008) and between 15-15.5 day hours by Grevstad (1999) based on conditions inducing oviposition during rearing. Individual variation in diapause induction at the same day length exposure shows genetic variation in the critical photoperiod within these source populations (Velarde et al. 2002, Bartelt et al. 2008). As in other insects, additional factors may modulate the photoperiod-based diapause response (Tauber et al. 1986).

Source populations in Germany were reported as primarily univoltine, with some oviposition from the summer generation before entering diapause (Blossey 1995). In the United States, 1 to 3 generations have been observed depending on location. The lifecycle model of Grevstad and Coop (2015) predicts that up to 4 generations are possible for *N. calmariensis* in the warmest parts of the new range based on available degree-days. However, its short-day diapause response would restrict it to fewer generations. Voltinism in the field, which may refer to either the completed or attempted (and not completed) number of generations, can be predicted with the assumptions that critical photoperiods approximate the responses to day length cues in natural settings and that day length provides the principal environmental cue. If day length was longer than the critical photoperiod during the stage at which *N. calmariensis* are sensitive, then most of the population would be reproductive.

We studied seven beetle populations that established in the states of California, Oregon, and Washington after their introduction in the 1990s (Figure 1). Environmental conditions at these sites vary with elevation (17-1,006 meters above sea level) and with latitude (39.41°-48.76°) (Table 1). For each location, we summarized the average annual season length in degree-days over 25 years (1994-2018) using Daymet daily gridded interpolations of weather station observations (Thornton et al. 1997, 2017). Grevstad and Coop’s (2015) degree-day model was used to predict average phenology of the F1 teneral adult eclosion, which varies with latitude and the day of year (Forsythe et al. 1995). We selected sites with established populations of beetles without recent transfers to our knowledge. The original releases in Washington used beetles from both northern and southern German sources, while those in Oregon only used beetles from the southern German source (Hight et al. 1995). California populations were introduced, often multiple times, from established populations in Oregon and Washington (M. Pitcairn personal communication) and could not establish south of 39° latitude (Pitcairn 2018). We summarize the known release history and translocations in Table 1, but other exchanges between populations that would alter their genetic variation and evolution are possible. In 2014, voltinism in the field was determined by the presence or absence of larvae in the second and third (where appropriate) generation after first confirming beetles were abundant early in the season. These surveys were timed to coincide with the predicted peak larval abundance rather than adults, because of the greater possibility for ambiguity about their generation of origin.

**Table 1:** Description of sites in the western USA with introduced *Neogalerucella calmariensis* populations.

| Site | Elevation<br>(meters<br>above<br>sea level) | Max<br>day<br>length<br>(hours)<sup>a</sup> | Avg<br>annual<br>degree-days<br>(SD)<sup>b</sup> | Average<br>F1 adult<br>emergence<br>prediction<sup>c</sup> | Population history (year introduced)<sup>d</sup> | Voltinism in the field<sup>e</sup> |
| --- | --- | --- | --- | --- | --- | --- |
| Bellingham, Washington<br>(48.76°, -122.48°) | 17 | 16.3 | 1067<br>(122) | July 20 | Introduced from Ephrata, WA<br>(unknown year in 1990s) | 1 generation |
| Ephrata, Washington<br>(47.16°, -119.66°) | 350 | 16.1 | 1677<br>(127) | June 24 | Original release from north and south<br>German sources (1992-93) | 3 generations |
| Yakima Training Center,<br>Washington (46.76°, -119.99°) | 204 | 16.0 | 1809<br>(131) | June 19 | Introduced from Ephrata, WA<br>(unknown year in 1990s). | Not observed, assumed similar<br>to nearby Ephrata, WA |
| Rickreall, Oregon<br>(44.98°, -123.27°) | 64 | 15.8 | 1366<br>(114) | July 9 | Original release from south German<br>source (1992) | 1 generation |
| Sutherlin, Oregon<br>(43.39°, -123.32°) | 153 | 15.6 | 1601<br>(143) | June 27 | Original release from south German<br>source (1992) | 2 generations |
| McArthur, California<br>(41.10°, -121.41°) | 1006 | 15.3 | 1559<br>(122) | July 1 | Introduced from Rickreall, OR and<br>vicinity (1998, 2000) & Ephrata, WA<br>(2002) | 2 generations |
| Palermo, California<br>(39.41°, -121.58°) | 35 | 15.1 | 2769<br>(175) | May 13 | Introduced from Rickreall, OR and<br>vicinity (1998-2001) & McArthur, CA<br>(2006). Release from Ephrata, WA<br>10km northwest of this site (2004-05)<br>could have dispersed here. | 3 generations |

<sup>a</sup>Maximum day length includes twilight until the sun is 1.5° below the horizon (using method of Forsythe et al. 1995).
<sup>b</sup>Degree-days are calculated with the triangle method with 10°C/30°C thresholds over 25 years (1994-2018) from Daymet daily gridded interpolations of weather station observations.
<sup>c</sup>Predicted sensitive stage emergence is based on 25 years (1994-2018) of results from a degree-day lifecycle model (Grevstad and Coop 2015).
<sup>d</sup>History is from Hight et al. (1995) for original releases in Washington and Oregon and from M. Pitcairn (personal communication) for transfers to California
<sup>e</sup>F. Grevstad and M. Pitcairn surveyed sites in 2014 to coincide with the expected timing of larval emergence in later generations.

**Figure 1:**
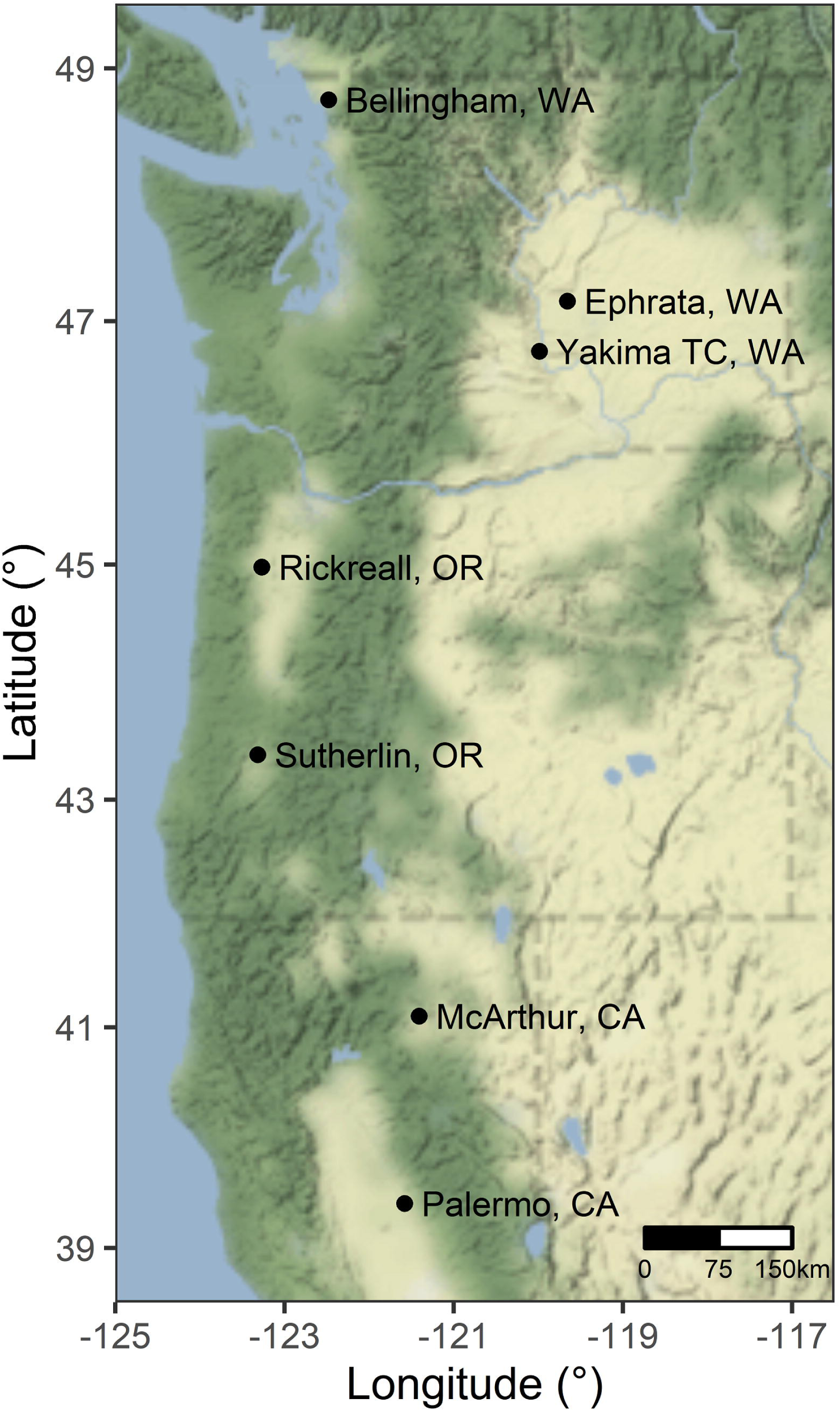
*Neogalerucella calmariensis* populations in the northwest USA in this study. Map tiles by Stamen Design (CC BY 3.0) with data by OpenStreetMap (ODbL) are plotted with the *ggmap* R package (Kahle and Wickham 2013).

### Experimental setup

We carried out two similar experiments (in 2014 and 2019) to compare the diapause response to photoperiod among *N. calmariensis* populations and one experiment (in 2018) to compare their rates of development from egg to adult. All experiments were performed in SG3-22 controlled environmental chambers (Hoffman Manufacturing, Inc.) equipped with six 32-watt 5000K tube fluorescent lights on the walls of each chamber. We reared populations in this controlled environment to attribute phenotypic variation to genetic differences between populations (Merilä and Hendry 2014). Beetles were collected from source populations in 2014 and 2018 for our lab colonies. At least one generation of rearing in a common environment preceded each experiment to control for maternal effects.

We grew *L. salicaria* host plants from seeds collected in Portland, Oregon (45.471°, −122.657°). Seedlings were transplanted to square 2.2-L plastic nursery pots with LA4PC potting soil (Sun Gro Horticulture H, Agawam, MA) and 5 grams of 14-14-14 slow-release fertilizer. Plants were maintained in standing water in a greenhouse with day/night temperatures of approximately 27/21°C and supplemental grow lights that turned on to extend the day length to 16 hours. We trimmed *L. salicaria* frequently to prevent flowering and to encourage growth of new shoots from axillary buds. When using cut leaves to feed *N. calmariensis*, we used new growth including shoot apical meristems and the first few sets of leaves.

The experiments differed in the method details as described below.

### Photoperiod-based diapause: 2014 experiment

*Neogalerucella calmariensis* eggs and early instar larvae were collected from established field populations in Palermo, California and Sutherlin, Oregon in early May and adults were collected near Ephrata and Bellingham, Washington in late May. We reared them in a greenhouse inside 13.5 × 13.5 × 24-inch mesh cages on potted *L. salicaria* until the next generation’s adults emerged. After the new adults started mating, we obtained same-age cohorts of eggs by placing 3 adult pairs on each of 12 caged *L. salicaria* (15-20 cm tall) for each population. This resulted in roughly 90 eggs per plant. When larvae started to hatch, whole plants were placed into growth chamber treatments (3 plants x 4 populations x 4 photoperiod treatments) and beetles developed to adults. The photoperiod treatments in chambers were 14.75/9.25, 15.25/8.75, 15.75/8.25, and 16.25/7.75 hours of light/dark (L/D) per day. To provide additional light for plant growth, two 24-watt 6500K tube fluorescent tube lights were suspended above the plants as a supplement to the built-in lights on the walls of the chamber. These were set to go on 15 minutes after and off 15 minutes before the built-in lights to emulate twilight. Chambers were maintained at 23°C constant temperature for day and night. Adults were removed from plants as they emerged and grouped in 150mm-diameter petri dishes in the chambers with cut leaves for the pre-oviposition or pre-diapause development time.

We assessed the developmental status (reproductive versus diapausing) in all adult females (556 individuals total) 10-14 days after their emergence, which was sufficient time for completion of pre-oviposition or pre-diapause development. Adults were placed individually into 50mm-diameter petri dishes with fresh leaves and moist filter paper for 48 hours. We assigned beetles as reproductive if oviposition occurred. Indeterminate behavior, such as feeding with no eggs, led to an extra 24-hour observation before classification. Diapause beetles had no feeding and were often hiding and inactive underneath the filter paper. The proportion of females that were reproductive for each treatment combination (pooled across plants) was the dependent variable for our analysis.

### Development time: 2018 experiment

To determine if development rates differed among populations of *N. calmariensis*, we carried out a separate experiment using the same environmental chambers. We intended this experiment to measure both diapause response and development timing. However, a high diapause rate across all photoperiods (at 21°C) meant that we were unable to obtain and compare diapause response curves. Therefore, we present it to compare development times between populations and test whether shorter photoperiods accelerate development.

We collected 50-100 adult beetles from two sites in Washington (Bellingham and Yakima Training Center) and two sites in Oregon (Rickreall and Sutherlin) between April 26 and May 8, 2018. Beetles from the two sites in California (McArthur and Palermo) were collected a month later. After rearing them for a generation in a greenhouse, we placed 30-35 beetles on individual, caged potted *L. salicaria* for 24 hours to establish same-age cohorts of eggs. After removing the adults, the plants with affixed eggs were placed in a constant 21°C growth chamber at 16.75/7.75 L/D hours. Shortly before hatching, eggs from the same cohort were distributed to 150mm-diameter petri dishes with fresh leaves lined with moist filter paper at a density of 30 to 50 eggs per dish. Each dish was randomly assigned to photoperiod treatments of 14.75/9.25, 15.75/8.25, 16.25/7.75, and 16.75/7.25 L/D hours (3 cohorts x 6 populations x 4 photoperiod treatments). We exchanged fresh leaves as needed and added 10 grams of potting soil substrate for pupation to each dish when the beetles reached the third instar. Adults were removed and counted every day as they emerged. In total, we tracked the development of 1808 beetles. In our statistical analysis, the duration from oviposition to adult eclosion for each individual was the dependent variable compared across populations.

### Photoperiod-based diapause: 2019 experiment

Diapausing beetles from the two shortest day length treatments in the 2018 experiment were stored outdoors in mesh cages with leaf litter from November 2018 through April 2019 in Corvallis, Oregon. We only used beetles from these two photoperiod treatments, which all went into diapause, so that we were not artificially selecting beetles based on their photoperiod response. These beetles emerged reproductive in the spring and their offspring entered this experiment. Experimental methods differed from the diapause experiment carried out in 2014 in that larvae were reared in 150mm-diameter petri dishes (approximately 30 individuals per dish), no supplemental lights were added to the chambers (used only built-in lights), and both males and females were assessed for diapause. We used the same rearing methods as the 2018 development time experiment, but with a higher constant temperature of 23°C and an extended range of photoperiod treatments compared to the 2014 and 2018 experiments. We assigned rearing dishes from each population to photoperiod treatments at 14.5/9.5, 15.0/9.0, 15.5/8.5, 16.0/8.0, 16.5/7.5, and 17.0/7.0 L/D hours (2 dishes x 6 populations x 6 photoperiod treatments). As they emerged each day, we removed and re-pooled teneral adults into group trays with fresh leaves for the pre-oviposition or pre-diapause development period. In the diapause behavior test (described earlier), 1,782 beetles were scored as reproductive female, reproductive male, or diapause (sex not determined) according to their oviposition, feeding, and activity. The proportion reproductive for each treatment combination pooled across rearing dishes was the dependent variable for our analysis.

### Statistical analysis

We performed all analysis and visualization with R 3.6.2 (R Core Team 2019). First, each year’s diapause data (2014 and 2019) were analyzed separately for beetle reproductive rates with generalized linear mixed effects models with a binomial error distribution and logit link using the *lme4* package (Bates et al. 2015). The fixed effects of population and photoperiod treatment (scaled continuous covariate) and their interaction predicted the proportion reproductive in each treatment combination, accounting for the total number of beetles in each group. As these groups contained beetles from the same population reared in the same growth chamber, they were assigned random intercepts to account for non-independence and allow for overdispersion in the residual variance (Bolker 2015). We removed the interaction term if non-significant (*p* > 0.05) by a Chi-square test. Population differences in responses (estimated marginal means) were compared using the *emmeans* package with post-hoc pairwise comparisons while adjusting for multiple comparisons using Tukey’s method (Lenth 2020). Populations’ critical photoperiods (=critical day lengths), the hours of light exposure at which half of the beetles are predicted to diapause, were derived with 95% confidence intervals using inverse prediction from the generalized linear mixed models (Venables and Ripley 2002).

Second, we tested for a latitudinal gradient in the diapause response by analyzing the 2014 and 2019 experiments in the same model. We changed the predictor variables in the generalized linear mixed models to shift the focus from populations’ critical photoperiods to the latitudinal cline in the trait. The fixed effects were photoperiod treatment, latitude (scaled), year of experiment, and all interactions. We removed interaction terms if non-significant (*p* > 0.05) by a Chi-square test. The two experiments differed in methods such that the results from their separate models (see above) had substantial differences in the slope of the diapause response to photoperiod treatments. By including year and its interactions with other variables in this model, we account for the potential different responses to photoperiod treatment across the latitudinal gradient that might be artifacts from the experiments’ methods. In addition to the random intercept for treatment combination to control for the non-independence of beetles reared in the same chamber, we included a random intercept for population because we are attempting to make inferences about the larger, unsampled set of beetle populations along the latitudinal gradient rather than just these seven populations (Bolker 2015). The predicted change in critical photoperiods across the latitudinal gradient was derived from this model with 95% confidence intervals using simulations from the model parameters and their uncertainty (Population Prediction Intervals in Bolker (2008)).

Third, we tested if development time was affected by population and photoperiod exposure with a generalized linear mixed effects model with a gamma error distribution and log link using the *lme4* package (Bates et al. 2015). The fixed effects of population and photoperiod treatment (scaled continuous covariate) and their interaction predicted the duration from oviposition to adult eclosion in days for each beetle. Rearing dishes, the experimental unit, with beetles from the same population and oviposition day, were assigned random intercepts to account for their non-independence. We removed the interaction term if non-significant (*p* > 0.05) by a Chi-square test. Population differences in responses (estimated marginal means) were compared using the *emmeans* package with post-hoc pairwise comparisons while adjusting for multiple comparisons using Tukey’s method (Lenth 2020).

## Results

Introduced populations have established at sites with average season lengths from 1067-2769 cumulative growing degree-days (base 10°C) and two months difference in the predicted eclosion of F1 adults that would be sensitive to the photoperiod cues that induce diapause. Surveys confirmed voltinism of 1-3 generations in the field in 2014 (Table 1).

### Photoperiod-based diapause

All populations exhibited a short-day diapause response where shorter photoperiod treatments induced diapause and longer photoperiod treatments resulted in reproduction (Table 2, Figure 2, Figure 3). Populations differed in their critical photoperiods by up to 0.7 hours in the 2014 experiment and 2.3 hours in the 2019 experiment (Table 2). There is no interaction between population and photoperiod response in either experiment, meaning that the slopes of the photoperiod response curves do not significantly differ across populations (Table 3).

**Table 2:** Photoperiod responses of *Neogalerucella calmariensis* populations in two experiments at constant 23°C temperatures.

| Predictors | 2014 Photoperiod response |  |  |  |  | 2019 Photoperiod response |  |  |  |  |
| --- | --- | --- | --- | --- | --- | --- | --- | --- | --- | --- |
|  | Beta | Std. error | *P* | CP<sup>a</sup><br>(95% CI) | Population differences<sup>b</sup> | Beta | Std. error | *P* | CP<br>(95% CI) | Population differences |
| Photoperiod:<br>light hours | 4.05 | 0.67 | <0.001 | - | - | 1.29 | 0.16 | <0.001 | - | - |
| Bellingham,<br>Washington | -2.20 | 0.58 | <0.001 | 16.1<br>(15.9-16.4) | A | -1.36 | 0.40 | <0.001 | 16.6<br>(16.0-17.1) | A |
| Ephrata,<br>Washington | -2.43 | 0.59 | <0.001 | 16.2<br>(16.0-16.4) | A | - | - | - | - | - |
| Yakima TC,<br>Washington | - | - | - | - | - | -0.29 | 0.41 | 0.488 | 15.9<br>(15.5-16.4) | AB |
| Rickreall,<br>Oregon | - | - | - | - | - | -1.76 | 0.41 | <0.001 | 16.8<br>(16.3-17.3) | A |
| Sutherlin,<br>Oregon | 0.40 | 0.52 | 0.443 | 15.6<br>(15.4-15.8) | B | 1.84 | 0.40 | <0.001 | 14.5<br>(14.0-15.1) | C |
| McArthur,<br>California | - | - | - | - | - | -0.34 | 0.37 | 0.365 | 15.9<br>(15.5-16.4) | AB |
| Palermo,<br>California | 0.86 | 0.49 | 0.080 | 15.5<br>(15.3-15.7) | B | 0.95 | 0.38 | 0.013 | 15.1<br>(14.6-15.6) | BC |

<sup>a</sup>Critical photoperiod (CP), or the light hours at which 50% would enter diapause, was estimated for each population and year with 95% confidence intervals by inverse predictions from the models.
<sup>b</sup>Populations with significant differences have different letters, which come from pairwise comparison tests with Tukey's method to adjust *P* values.

**Table 3:** ANOVA for tests of photoperiod and population effects on diapause proportion and development time.

|  | 2014 Diapause |  |  | 2019 Diapause |  |  | 2018 Development time |  |  |
| --- | --- | --- | --- | --- | --- | --- | --- | --- | --- |
| Fixed effects | Chi square | df | P | Chi square | df | P | Chi square | df | P |
| Photoperiod (light hours)<sup>a</sup> | 36.6 | 1 | <0.001 | 82.7 | 1 | <0.001 | 0.0396 | 1 | 0.842 |
| Population | 25.8 | 3 | <0.001 | 68.7 | 5 | <0.001 | 14.8 | 5 | 0.0113 |
| Photoperiod x Population | 1.41 | 3 | 0.703 | 6.53 | 5 | 0.258 | 9.62 | 5 | 0.0869 |
| Random effects | Groups | SD |  | Groups | SD |  | Groups | SD |  |
| Intercept<sup>b</sup> | 16 | 0.602 |  | 36 | 0.708 |  | 71 | 0.0135 |  |

Type II sum of squares (Wald chi square tests) from generalized linear mixed models.
<sup>a</sup>Photoperiod is modeled as a continuous variable and had different growth chamber treatments in different experiments.
<sup>b</sup>Random intercepts in were based on treatment group for diapause models and rearing dish for development time model. Standard deviation (SD) shows between-group variation.

**Figure 2:**
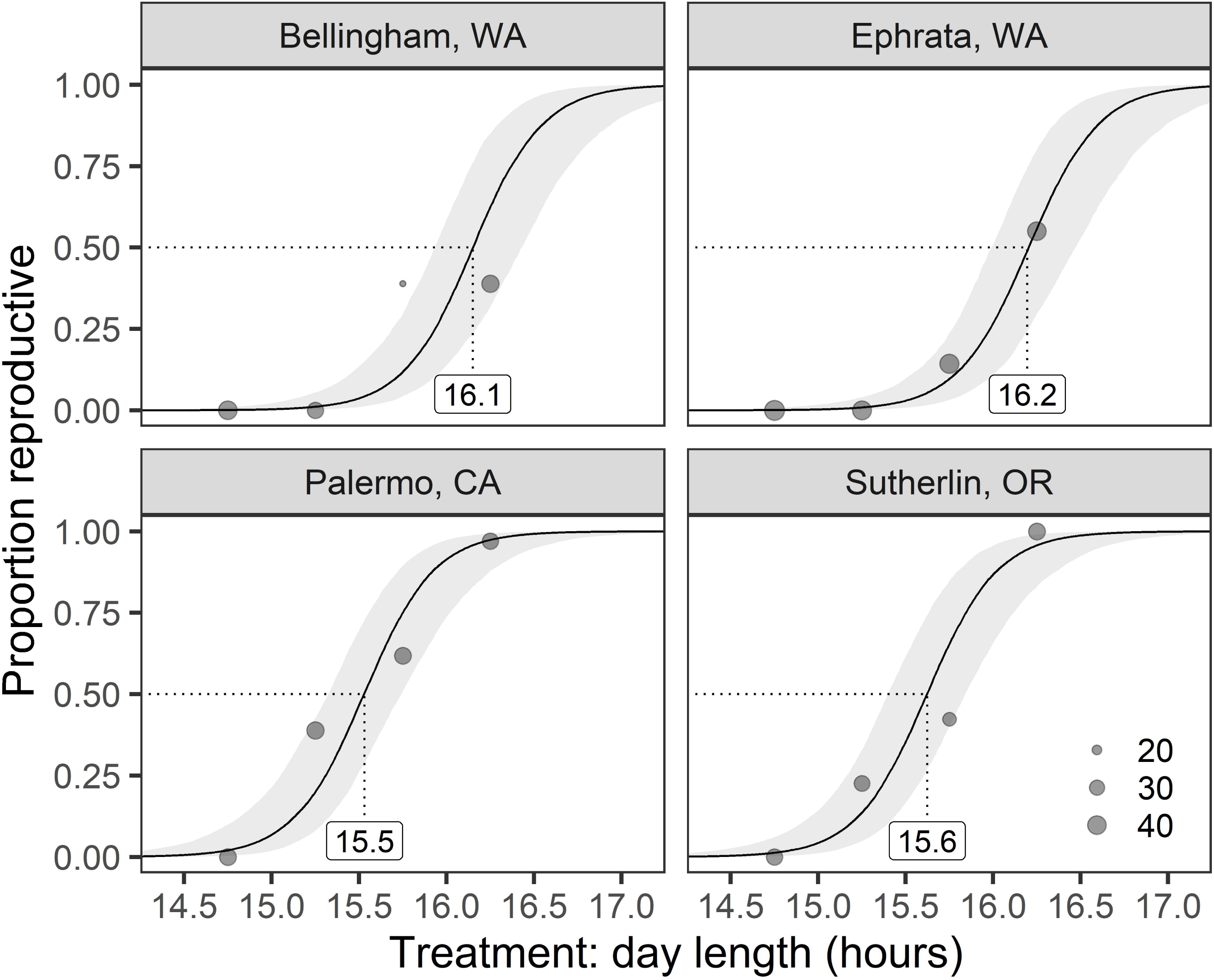
Photoperiod responses diverged between northern and southern populations in 2014. Each data point shows the proportion of female beetles that were reproductive for each treatment combination, with size scaled by number of beetles, for four populations (panels) and four photoperiod treatments (x-axis). Solid lines and 95% confidence intervals show generalized linear mixed model predictions. The critical photoperiods, at which 50% choose to diapause, are labeled below the dotted lines.

**Figure 3:**
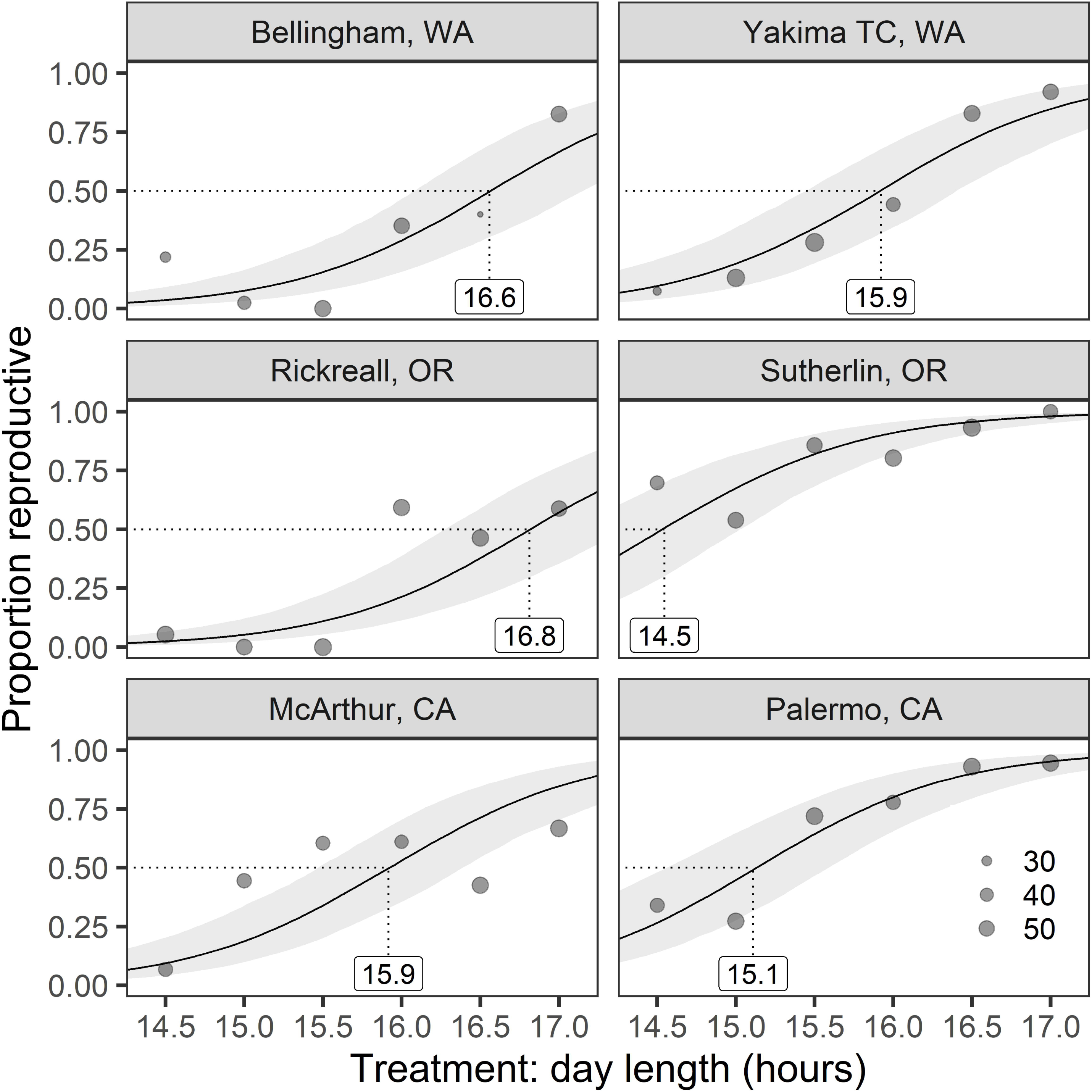
Photoperiod responses diverged between populations in 2019. Each data point shows the proportion of male and female beetles that were reproductive for each treatment combination, with size scaled by number of beetles, for six populations (panels) and six photoperiod treatments (x-axis). Solid lines and 95% confidence intervals show generalized linear mixed model predictions. The critical photoperiods, at which 50% choose to diapause, are labeled below the dotted lines.

Populations had a lower proportion reproductive, regardless of photoperiod, at higher latitudes (β = −0.918, SE = 0.393, *P* = 0.0196, Table 4). The 2019 experiment had marginally higher proportions reproductive (β = 0.747, SE = 0.414, *P* = 0.0709, Table 4) and a significant interaction with the photoperiod response (β = −2.97, SE = 0.643, *P* = <0.0001, Table 4). This interaction is visualized by the steeper slopes of the photoperiod response in 2014 (Figure 2) compared to 2019 (Figure 3). From this model, critical photoperiods increased by 15.7 (CI 13.9-17.4) minutes per 5° latitude in the 2014 experiment and 51.0 (CI 45.0-56.9) minutes per 5° latitude in the 2019 experiment (Figure 4).

**Table 4:**
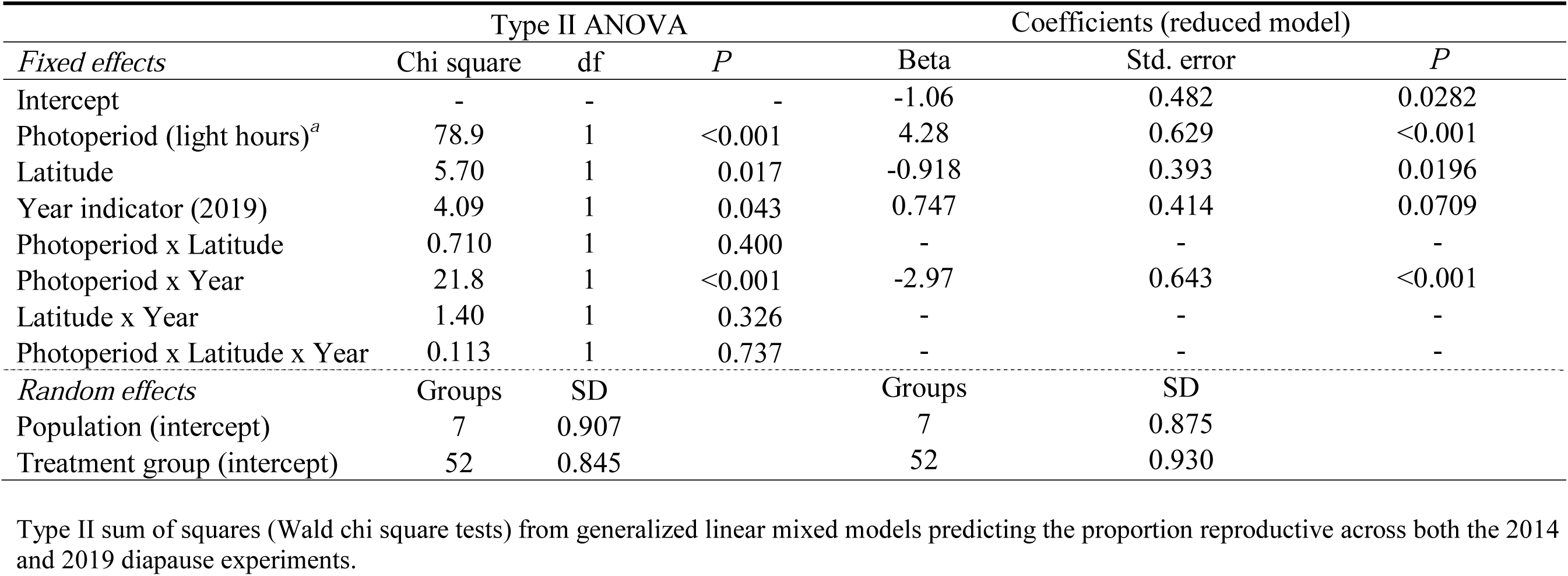
Test of latitudinal gradient in photoperiod effects on diapause proportion over two experiments.

**Figure 4:**
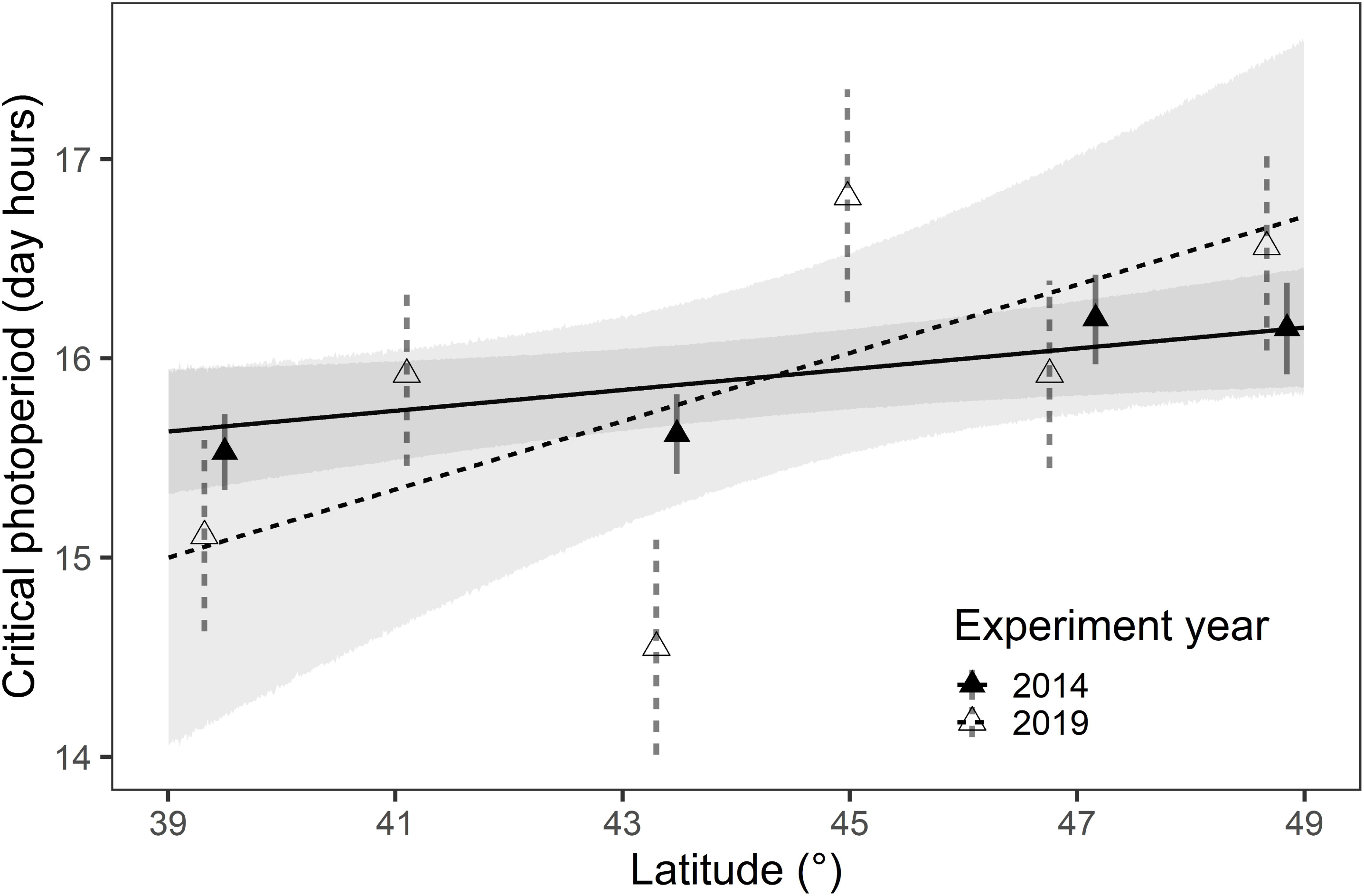
Latitudinal clines in critical photoperiod with estimated population traits. Lines shows the modeled change in critical photoperiod with latitude, estimated for the 2014 (solid) and 2019 (dashed) experiments. Confidence intervals (95%) for each line are shaded gray. Note that the lines do not significantly differ in slope (i.e. no latitude x year interaction, Table 4). Triangles show the critical photoperiods and 95% confidence intervals estimated for populations in separate models (corresponding to Figures 2 and 3). The x-axis values for the three sites with repeat measures were offset by 0.1° to avoid overlap.

However, the six populations tested in 2019 do not have a monotonic increase in critical photoperiod with latitude (Figure 4, Table 2). For example, the two central populations (Rickreall and Sutherlin, Oregon) have the longest (16.8, CI 16.3-17.3) and shortest (14.5, CI 14.0-15.1) critical photoperiods, respectively (3.87 difference, SE = 0.551, *P* < 0.001 on logit scale). Populations from McArthur, California and Yakima Training Center, Washington have similar critical photoperiods in spite of their separation of 5.7° latitude (0.00679 difference, SE = 0.502, *P* > 0.999).

### Development time

Development time averaged 33.5 days and did not change with photoperiod treatments (Table 3, Figure 5). Populations had similar development times with the exception that the Rickreall, Oregon population required an additional day of development compared to the Sutherlin, Oregon population (0.0419 difference on log scale, SE = 0.0127, *P* = 0.0122) and the Yakima Training Center, Washington population (0.0364 difference on log scale, SE = 0.0127, *P* = 0.0471). There was no interaction between population and photoperiod for development time (Table 3).

**Figure 5:**
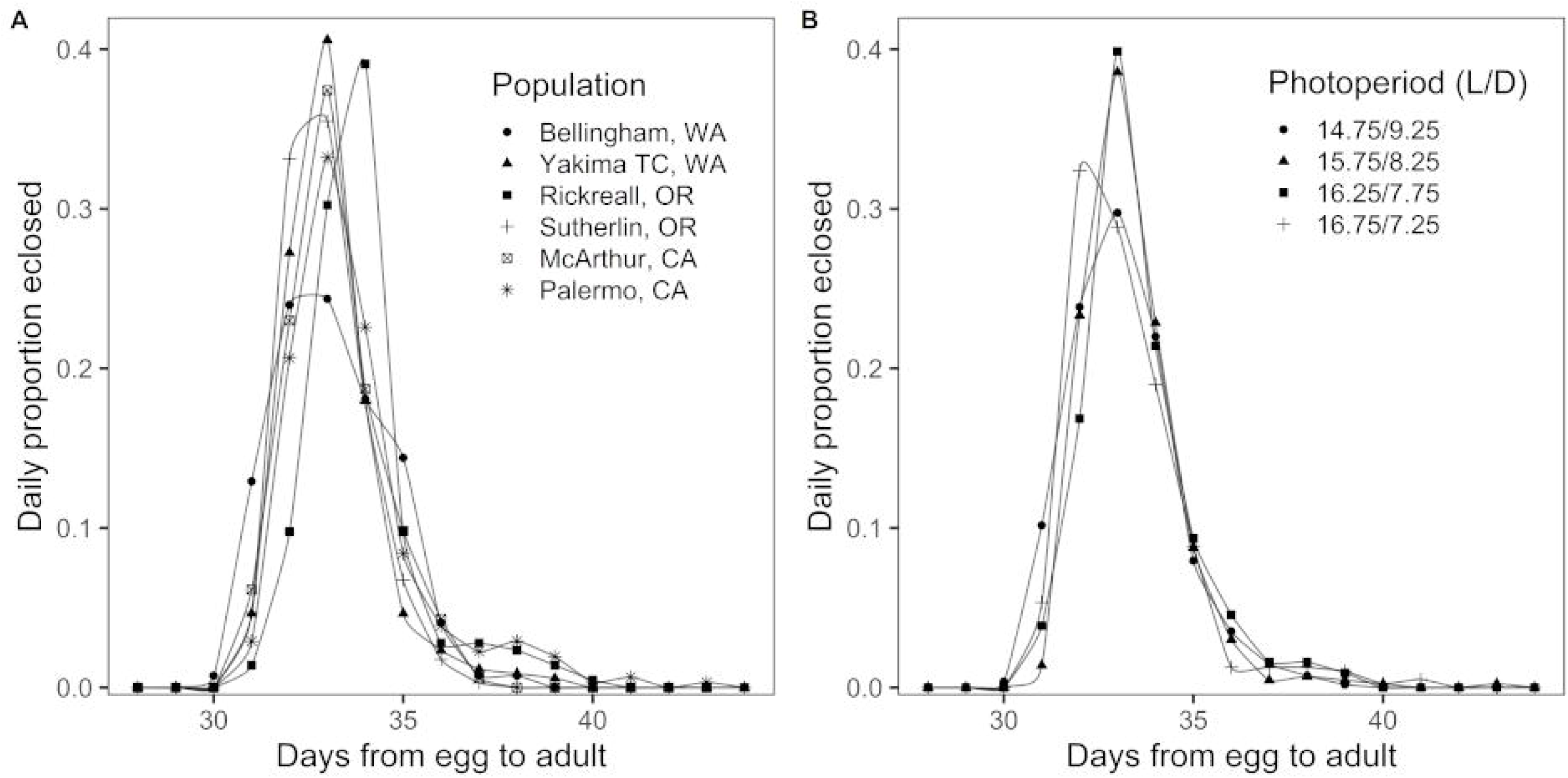
Main effects of population and photoperiod on development time at constant 21°C. Points represent daily observations of the proportion of the total number of beetles that eclosed in each group. Development time is similar for most populations, but the Rickreall, OR population took longer than the populations in Sutherlin, OR and Yakima, WA (A). Photoperiod treatments had no effect on development time in the four growth chambers (B).

## Discussion

Nearly three decades after their introduction, populations of the introduced leaf beetle *Neogalerucella calmariensis* have genetically diverged in their diapause response to photoperiod across 9.4° of latitude in the western USA. Despite a genetic bottleneck at the time of their introduction from Germany in the 1990’s, the measured critical photoperiods, where 50% of the population enters diapause, range from 14.5 to 16.8 hours of daylight, greatly extending the estimated 15 to 15.5 hour critical photoperiod in the original population. The measured critical photoperiods roughly correlate with latitude, with longer critical photoperiods found in the north where summer days are longer and where conditions favorable for reproduction tend to end earlier in the year (Figure 4). Critical photoperiods were shorter in the south where summer day lengths are shorter and conditions usually remain favorable later into the autumn.

The evolved geographic variation resembles the photoperiod response clines commonly seen in native or long established insects. Typical latitudinal gradients for critical photoperiods in locally-adapted insects in temperate zones range from 60-90 minutes per 5° latitude in classic experiments (Danilevskii 1961) or 48 minutes per 5° latitude in a recent meta-analysis (Joschinski and Bonte 2020). For the studied *N. calmariensis* populations, we estimated critical photoperiod clines of 15.7 minutes and 51.0 minutes per 5° latitude in 2014 and 2019 experiments, respectively (Figure 4). The lower rates of change with latitude found in this study may reflect incomplete adaptation in this introduced insect, or may be a result of the particular locations sampled (see below). If we assume that the latitudinal clines in native insects represent adaptations to the local seasonal environment, then we can say that *N. calmariensis* has evolved in an adaptive direction. However, reciprocal transplant experiments would be needed to confirm that these changes in photoperiod responses are local adaptations that increase fitness (Kawecki and Ebert 2004, Merilä and Hendry 2014, Tsai et al. 2020).

We found that development rates remain similar among populations with the exception of a slower development rate, by one day, in the Rickreall, Oregon population (Figure 5). The small difference, in comparison to the wider variation found in critical photoperiods, suggests that photoperiod response plays a relatively larger role in adapting to local seasonal regimes (Bradshaw and Holzapfel 2007). It is intriguing that this population with slower development also has a longer critical photoperiod than its neighboring populations. Field observations from recent years confirm a single generation at this site, whereas the Sutherlin, Oregon population just 1.6° latitude to the south and unsampled populations within 0.5° latitude to the north have 2 generations (Table 1, F. Grevstad personal observations). It is unclear why this population should be limited to one generation given sufficient degree days for two generations and plants that remain green until early fall. Relaxation of selection for faster development may be a result of the predominantly univoltine lifecycle at this site. Other insects have evolved nonlinear “stepped” or “sawtooth” clines in developmental traits across latitudes in tandem with locally-adapted photoperiod responses, which has the added effect of limiting attempted generations in places where season length is only marginally supportive of increased voltinism (Roff 1980, Tauber and Tauber 1981, Levy et al. 2015).

Measured critical photoperiods are useful in combination with developmental degree-day requirements for predicting voltinism at a given location over time (Beck and Apple 1961, Tobin et al. 2008, Kerr et al. 2020). The relationship of a population’s mean critical photoperiod to modeled voltinism for a given climate and latitude has been covered in detail in Grevstad and Coop (2015). In brief, a degree-day phenology model predicts when the photoperiod-sensitive stage emerges, and if the day length on that date for the latitude is longer than the critical photoperiod then the population is reproductive and attempts another generation. Voltinism can shift (sometimes dramatically) when an insect is moved to a new climate or latitude and with annual variation in phenology of the sensitive stage. However, laboratory-measured critical photoperiods may differ that those used in the field, where multiple cues interact and conditions vary from rearing experiments. For example, compared to the abrupt transition between light and dark in the lab, we do not know what proportion of the twilight period *N. calmariensis* perceives as part of the day length for comparison against the critical photoperiod. Where this has been studied in other insects, the portion of twilight sensed as part of the photoperiod appears be quite variable among insect species (Menzel 1979) and may even vary asymmetrically between dawn and dusk (Takeda and Masaki 1979). Other laboratory conditions, such as constant temperatures, may also change the estimated critical photoperiods. Our 2018 experiment anecdotally had much lower reproductive rates (and presumably higher critical photoperiods) after rearing at 2°C cooler temperatures. In other beetles, rearing at constant temperatures increased the measured critical photoperiod by 15 minutes compared to a fluctuating regime with the same mean (Bean et al. 2007). Now that our study has demonstrated genetic differences in a simplified experimental setting, field validation would help gauge how diapause initiation changes in the presence of twilight, temperature fluctuations, density-dependence, and host plant defenses.

The estimated critical photoperiods vary substantially between experiments and about the latitudinal trend. We cannot rule out that the photoperiod response evolved over a few years (Bean et al. 2012, Urbanski et al. 2012) or even changes annually with strong fluctuating selection (Bell 2010). However, methodological differences likely caused variation between years, such as the higher precision of critical photoperiods estimated in 2014. The use of whole potted plants for rearing in the 2014 experiment could have provided beetles with additional cues from the host plant to induce a stronger diapause response (Izzo et al. 2014). The inclusion of males in the sample in 2019, but not 2014, may have led to increased standard deviation in the critical photoperiod (Table 2) and the marginally significant higher proportion of reproduction beetles (Table 4). There are also climatic and phenological reasons why populations’ critical photoperiods vary from the latitudinal trend. In the western United States, local climates are influenced by complex topography relative to the eastern United States. The sawtooth pattern in critical photoperiods estimated in 2019 (Figure 4) correspond to differences in average degree-days and sensitive stage phenology at the sampled sites (Table 1). Local climate determines the number of generations that are possible in a location and the evolved critical photoperiod may vary over short distances at similar latitudes (Lindestad et al. 2019). Insect phenology models that incorporate demography along with photoperiod-based diapause can estimate annual fitness for a particular critical photoperiod (Kerr et al. 2020), and extending these models across years could answer whether a critical photoperiod has reached an optimum for a population’s environment (similar to trait models in Kivelä et al. (2013)).

The photoperiod response in *N. calmariensis* appears to be quite variable among individuals, with a mix of reproductive and diapausing beetles in the same treatment groups (Figures 2 and 3). While less individual variation, and a steeper slope in the photoperiod response curve, may show stronger selection on the critical photoperiod (Tauber et al. 1986), some insect species show substantial individual variation with flatter slopes (Joschinski and Bonte 2020). In the field, this variability can translate into partial generations toward the end of the season, where a portion of adults within the same generation enters diapause while the rest go on to reproduce. Two years of observations suggest that this commonly occurs in colonies reared outdoors in Corvallis, OR (T. Wepprich personal observation). The extent to which this occurs in natural populations of *N. calmariensis* would be worth further study. Higher individual variation, or a weaker diapause response to photoperiod cues, may facilitate insect invasions to new locations (Reznik et al. 2015) and may be a trait to screen in potential biological control agents.

Based on this study, there was sufficient genetic variation from two source populations to evolve divergent photoperiod responses across a range of environmental conditions within 27 years. The evolution of *N. calmariensis*, as well as other biological control agents (Bean et al. 2012, McEvoy et al. 2012, Szucs et al. 2012), suggests the importance of season length as a selective force over ecological timescales. Screening for developmental traits or environmental factors that predict successful establishment and population growth of biological control agents would advance our ability to manage planned introductions (Zalucki and van Klinken 2006, Abram and Moffat 2018). More targeted matching of observed photoperiod responses in the native range combined with lifecycle simulations of growing season lengths in the introduced range may reduce the length of time or number of introductions needed to establish robust populations (Grevstad and Coop 2015, Pitcairn 2018). Beyond the field of biological control, evolution to changing seasonality is the predominant genetic response to rapid climate change in natural populations (Bradshaw and Holzapfel 2006) and photoperiodism will likely determine range shifts across diverse taxa (Saikkonen et al. 2012). Quantifying changing photoperiod responses in introduced populations provides examples of how quickly and to what extent evolution can track anthropogenic warming, with the changes in both season length and timing of environmental cues it brings.

## Acknowledgments

We thank Lisa Weigel at Yakima Training Center and Mike Pitcairn and Baldo Villegas at the California Department of Food and Agriculture for support with site selection and collections. We thank Brittany Barker, Len Coop, Katarina Lunde, Monte Mattson, and Peter McEvoy for feedback on manuscript drafts. FG received funding from the DOD Strategic Environmental Research and Development Program (US Army Corps of Engineers, Contract No. W912HQ-17-C-0051) and the USDA Forest Service cooperative agreement 10-CA-11420004-255.

## Notes

### Competing Interest Statement

The authors have declared no competing interest.

### Summary of Updates

Thanks to the suggestions of 3 anonymous peer reviewers, we made the following major changes: 1.We expanded the methods as suggested with more details on the lifecycle, site history, and experimental setup. For some details, we validated experimental protocols with our notes and changed the descriptions if warranted. 2.For additional context about voltinism in the field, we added results from surveys conducted in 2014 at 6 of the sites to verify reproduction of later generations. 3.We added a new analysis (and corresponding Figure 4 and Table 4) combining the 2014 and 2019 diapause experiments to directly test for a latitudinal cline in critical photoperiod. We take care to emphasis that these two experiments differ in their methods, controlled for in this new analysis with interaction terms, and that the results do not necessarily indicate evolution over the few years between experiments. 4.We reevaluated our choices for grouping non-independent experimental observations in the analysis. In the diapause experiments, we moved beetles from rearing groups into new group dishes as they emerged for the pre-oviposition period. Due to this mixing, the best way to account for similar rearing conditions is by pooling individuals by treatment combination rather than rearing dish (as done in the original analysis). This choice did not change the significance or interpretation of the fixed effects in any of the results. 5.We reworked the discussion completely to include the new analysis of the latitudinal gradient in critical photoperiods and to explain our expectations for how our experimental results would apply to voltinism in the field.

